# Proteomics reveals synergy in biomass conversion between fungal enzymes and inorganic Fenton chemistry in leaf-cutting ant colonies

**DOI:** 10.1101/2020.08.11.239541

**Authors:** Morten Schiøtt, Jacobus J. Boomsma

**Affiliations:** Centre for Social Evolution, Department of Biology, University of Copenhagen, Universitetsparken 15, DK-2100 Copenhagen, Denmark

## Abstract

The herbivorous symbiosis between leaf-cutting ants and fungal cultivars processes biomass via ant fecal fluid mixed with munched plant substrate before fungal degradation. Here we present a full proteome of the fecal fluid of *Acromyrmex* leaf-cutting ants, showing that most proteins function as biomass degrading enzymes and that ca. 80% are produced by the fungal cultivar and ingested, but not digested, by the ants. Hydrogen peroxide producing oxidoreductases were remarkably common in the fecal proteome, inspiring us to test a scenario in which hydrogen peroxide reacts with iron in the fecal fluid to form reactive oxygen radicals after which oxidized iron is reduced by other fecal-fluid enzymes. Our biochemical assays confirmed that these cyclical Fenton reactions do indeed take place in special substrate pellets, presumably to degrade recalcitrant lignocellulose. This implies that the symbiosis manages a combination of chemical and enzymatic degradation, an achievement that surpasses current human bioconversion technology.

---

Mutualistic mergers of simpler biological entities into organizationally complex symbioses have been key steps in the evolution of advanced synergistic forms of life, but understanding the origins and secondary elaborations of such natural cooperative systems remains a major challenge for evolutionary biology (1, 2). Physiological complementarity may be a main driver to maintain symbiotic associations, but new adaptations are also expected to jointly exploit the full potential of symbiosis (3, 4). The leaf-cutting ants belong to the genera *Acromyrmex* and *Atta* and form the phylogenetic crown group of the attine fungus-growing ants, which all live in mutualistic symbiosis with fungus-garden symbionts (5). While the basal branches of the attine clade farm a considerable diversity of mostly leucocoprinaceous fungi, normally in underground chambers (6), the leaf-cutting ants almost exclusively farm what has been described as the highly variable cultivar species *Leucocoprinus gongylophorus* (7). The ants cut leaf fragments from the canopy and understory and bring them back to their nest to be processed as growth substrate for the cultivar, which involves munching forage material into smaller pieces and mixing this pulp with droplets of fecal fluid before depositing the mixture as manure on the actively growing parts of the garden (8).

The deep ancestors of the attine ants and their fungal cultivars were hunter-gatherers and saprotrophs, respectively, which implies that none of them were adapted for efficiently degrading fresh plant material. Neither was there any need initially, because all phylogenetically basal attine ants provision their gardens with dead plant debris. There is a striking difference between the small colonies of these basal attine ants and the large colonies of leaf-cutting ants which are conspicuous functional herbivores with a large ecological footprint (9, 10). However, most of the leaves that the ants harvest are heavily protected against herbivory by recalcitrant cell walls and toxic substances (11), defences that needed to be overcome for obtaining net nutrition. In recent decades, a substantial amount of research has elucidated the selection forces that shaped the functional herbivory niche of the leaf-cutting ants. It was shown that: 1. The ‘higher’ attine ants to which the leaf-cutting ants belong rear specialized, gongylidia-producing cultivars that lost their free-living relatives and could thus begin to co-evolve with the ants (7); 2. The leaf-cutting ant cultivar *Leucocoprinus gongylophorus* evolved to be obligatorily multinucleate and polyploid, which may well have allowed higher crop productivity (12); 3. Leaf-cutting ant queens evolved to be multiply inseminated, in contrast to all phylogenetically more basal attines studied, which meant that not only the garden symbiont but also the farming ant families became genetic chimeras (13); 4. The leaf-cutting ants evolved distinct division of labor within the worker caste, with small, medium and large workers and specialized soldiers (the latter only in *Atta*) (8); and 5. The crown group of the attine ants underwent significant changes in their gut microbiota that likely facilitated the conversion efficiency of fungal diet substrate into ant biomass (14, 15).

While these advances have significantly enriched our understanding of the spectacular biology of the attine ants in general and the leaf-cutting ants in particular, very few studies have addressed the molecular synergy mechanisms that were decisive for integrating the complementary innovations of the ants and their cultivars. Recent genome sequencing showed that cultivars likely changed their chitinase processing abilities in co-evolution with the ants’ labial glands (16), but the clearest and most detailed evidence of functional physiological complementarity has been found for the digestive enzymes of the mutualistic farming symbiosis. These insights were initiated by pioneering work by Michael M. Martin (17–20), which suggested that proteases in the fecal fluid of *Atta* leaf-cutting ants had actually been produced by the fungal symbiont and passed unharmed through the ant guts to mediate protein degradation in the plant forage material when it was mixed with fecal fluid. Our previous work not only confirmed this hypothesis (21), but also expanded the list of cultivar-derived fecal fluid proteins to include pectinases, hemicellulases and laccases (22–25), while showing that all these enzymes are produced in the specialized gongylidia of the fungal cultivar that the ants ingest as their almost exclusive food source (21, 23–25).

This work also found a number of puzzling deficiencies in the form of decomposition elements that were missing. First, the degradation of leaf fragments in the fungus garden proceeds very efficiently, as ant-produced pellets of munched leaf material deposited on top of the garden turn black within a few hours to then be spread out over the garden by the ants. Likewise, the fungal cultivar has lost a major ligninase domain (16), begging the question how these recalcitrant cell wall components are breached to allow fungal hyphae access to the nutritious interior of live plant cells. To help resolve this puzzle, the present study focused on obtaining a full proteome of the fecal fluid of the leaf-cutting ant *Acromyrmex echinatior*, extending the earlier partial proteome from 33 to 101 identified proteins (23). This revealed a surprisingly high proportion of oxidoreductases, which made us conjecture that the mutualistic ant and fungal partners might jointly realize aggressive biomass conversion via the inorganic Fenton reaction, which produces hydroxyl radicals that can break down lignocellulose. We thus set up a series of experiments to investigate whether Fenton chemistry indeed takes place when fecal fluid interacts with munched leaf pulp concentrated in the 3 mm pellets that the ants construct. We confirmed our basic hypothesis and also found other enzymatic and iron components of the fecal fluid participating in the recycling of Fenton chemistry reactants. The synergistic process of enzymatic and inorganic chemistry appears to sustain a continuous production of hydroxyl radicals until the ants terminate the process by dismantling their temporary Fenton bioconversion pellets to offer the pre-digested plant substrate to the fungal hyphae and seemingly without any collateral damage to ant or fungal tissues due to uncontrolled hydroxyl radicals.

## Results

### Characterization of the fecal fluid proteome

Our proteomic analysis of worker fecal fluid from four different *A. echinatior* colonies identified 174 proteins (Supplementary Table S1), which showed substantial overlap among the pooled colony-specific samples (Fig 1A). To avoid including spurious proteins, we restricted our analyses to a shortlist of 101 proteins present in at least three of the four colony samples, and we were able to annotate 91 (90%) of these proteins (Table 1). This core proteome consisted mainly of degradation enzymes belonging to five major functional categories: oxidoreductases, proteolytic enzymes, carbohydrate active enzymes (cazymes), phosphate-liberating enzymes, and lipid degrading enzymes (lipidolytic enzymes). Of the ten remaining proteins, only three could not be functionally annotated. The vast majority (86) of the core fecal proteome originated from the fungal symbiont, with only 15 being encoded by the ant genome. Based on the number of spectra obtained for each identified protein, MaxQuant produced an approximate measure of relative abundance of each protein (LFQ intensity) (Table 1), which revealed that lipidolytic enzymes constitute only a very small fraction and that the “other” category was of intermediate size. The four remaining categories had summed LFQ intensities of the same order of magnitude (Fig 1B) and two specific proteins stood out as being exceptionally abundant (Supplementary Table S1), the Alkaline phosphatase (FP075) and the previously described Laccase LgLcc1 (FP097) (24), which jointly made up almost a quarter of the total fecal protein mass.

**Table 1.**
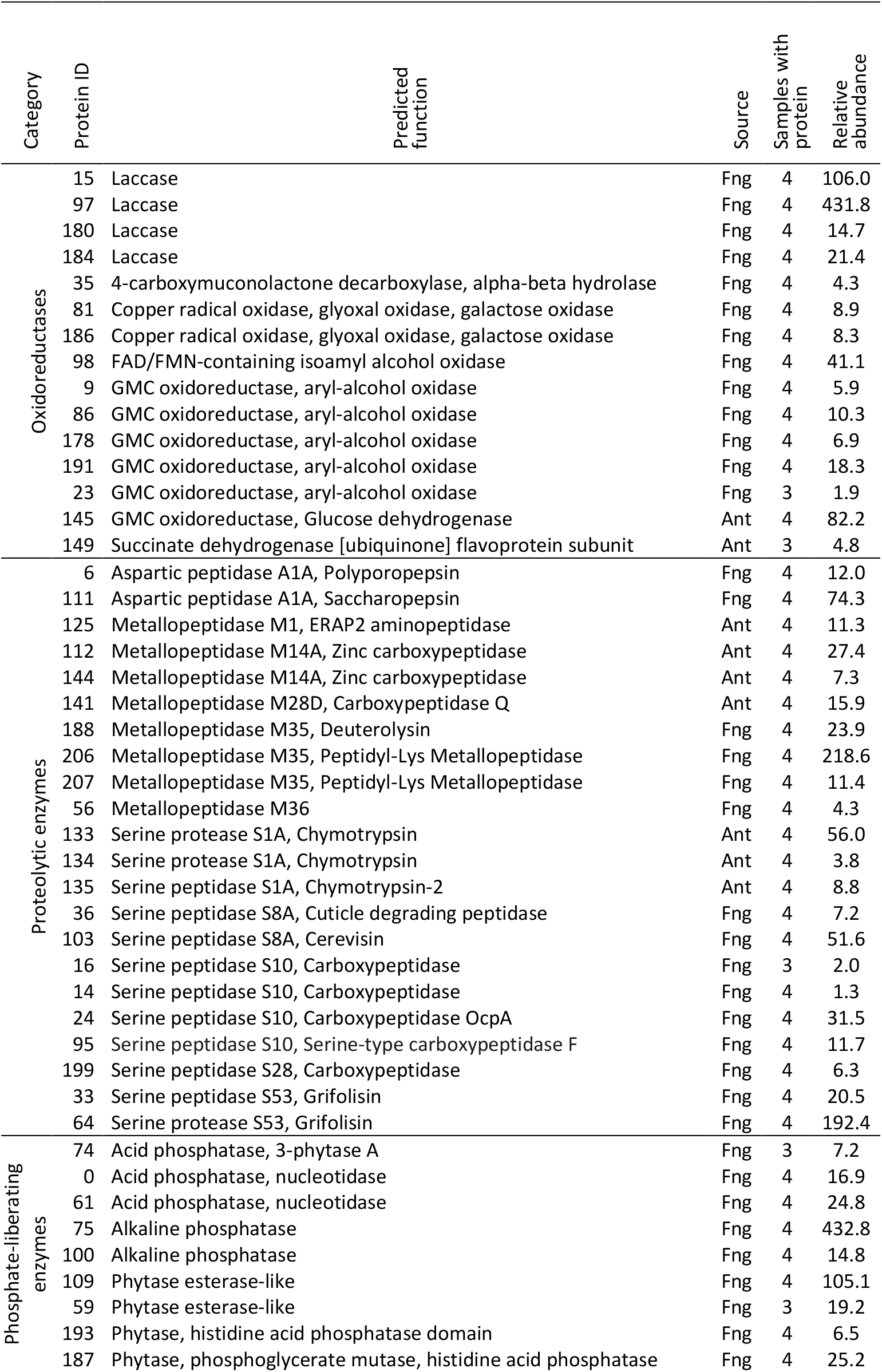
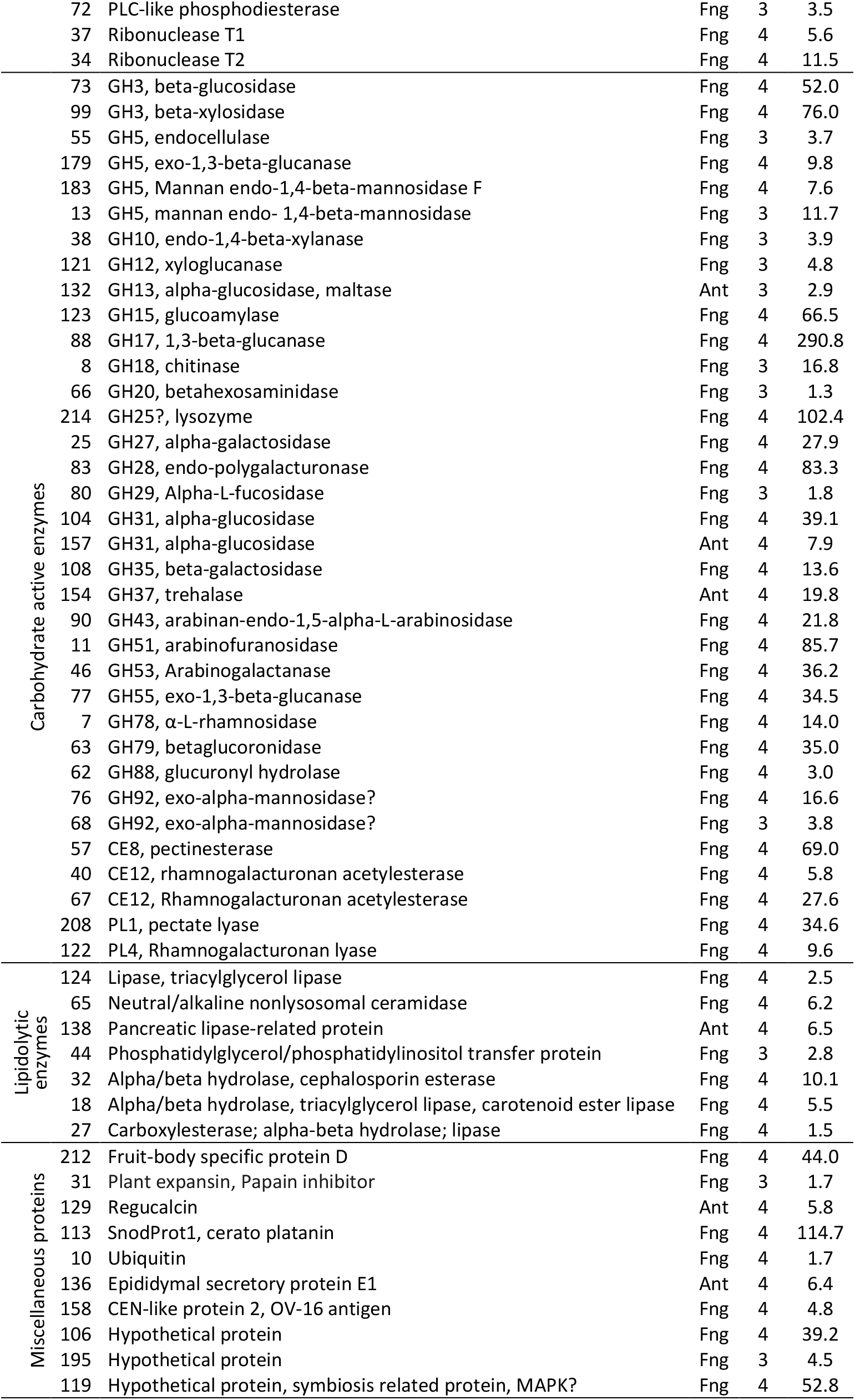
Fecal fluid proteins that were found in 3 or 4 of the examined colony samples (cf. Figure 1), and thus likely to belong to the core fecal fluid proteome. For each protein the predicted function, its fungal or ant origin, the number of samples containing the protein, the relative abundance of the protein, and an identifier number are listed. Proteins were assigned to one of six functional categories (cf Figure 1). Questionmarks indicate that annotations were inconclusive.

**Figure 1.**
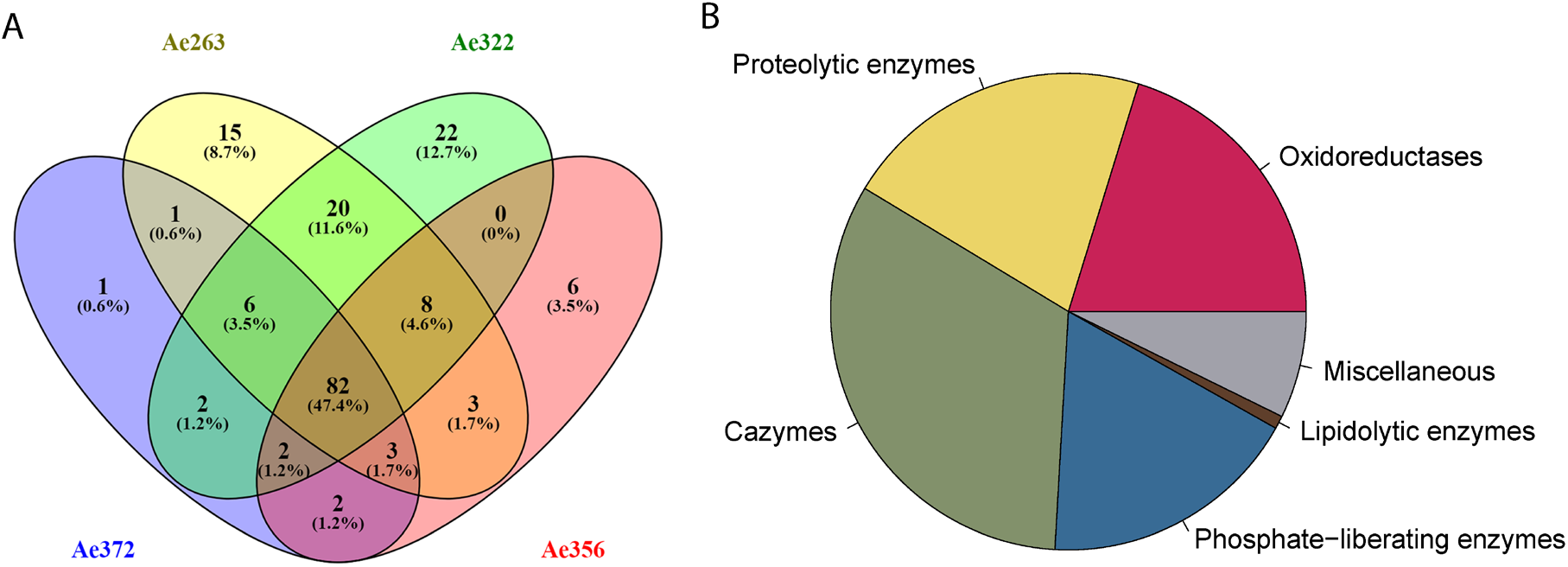
Statistics of the fecal fluid proteome of *Acromyrmex echinatior*. (**A**) Venn diagrams, constructed using the web application Venny 2.1; http://bioinfogp.cnb.csic.es/tools/venny/index.html, showing the overlap of protein profiles identified in the four fecal fluid samples obtained from colonies Ae263, Ae322, Ae356 and Ae372. (**B**) Pie chart showing the abundances of proteins across colonies assigned to six functional categories based on the LFQ values provided by MaxQuant.

Three other proteins also showed remarkably high abundance levels, a 1,3-beta-glucanase (FP088), a Metallopeptidase M35 (FP206), and a Serine protease S53 (FP064), which implies that the five top abundant proteins contributed more than 40% of the total fecal protein mass. These results confirmed our previous findings that a large proportion of the fecal fluid proteins consists of proteases (21), pectinases (23), hemicellulases (25), and laccases (24), but we now also identified several phosphate-liberating and lipidolytic enzymes. Just like nitrogen and carbon, phosphorus is a major macronutrient required for many biological processes. Enzymatic degradation of organic matter to liberate C, N and P is known to take place in a stoichiometrically balanced way (26), so it is not surprising that the fecal fluid also contains a range of enzymes able to release phosphorus from various substrates. That alkaline phosphatase (FP75) had the highest abundance of all fecal proteins underlines the importance of phosphorus mineralization. Several enzymes able to release phosphorus from phytin were also present. This is a storage compound for phosphorus in plant seeds, but also occurs in pollen and vegetative tissues (27). Laboratory colonies of *A. echinatior* were occasionally fed with dry rice grains, but in nature these ants are not known to forage on seeds to any significant degree (28). However, they do harvest flowers, whose pollen may be a source of phytin. Finally, the fecal fluid contained ribonucleases that will degrade RNA, another major source of phosphorus. The fecal lipidolytic enzymes had low abundances overall. They probably serve to degrade plant cell membranes to liberate cytosolic compounds which would release nutritious assimilatory lipids and phosphate present in cell membrane phospholipids.

### The complementary components for Fenton chemistry

Five of the identified oxidoreductases were annotated as fungal GMC oxidoreductases. A subgroup of these enzymes are the aryl alcohol oxidases that are believed to play an important role in lignin degradation by brown-rot fungi (29). These fungi produce aromatic alcohols, especially veratryl alcohol, which is subsequently oxidized by aryl alcohol oxidases to produce hydrogen peroxide (H_2_O_2_). The hydrogen peroxide is subsequently used in a Fenton reaction in which reduced iron (Fe^2+^) reacts with hydrogen peroxide to produce hydroxyl radicals that can diffuse into the lignocellulose substrate and break the chemical bonds responsible for the rigid structure of lignocellulose (Fig 2). This in turn will loosen the cell walls and allow access for more bulky degradation enzymes to target specific components of the plant cell wall. Unless the oxidized iron (Fe^3+^) is recycled by reduction to Fe^2+^, the production of reactive oxygen species will end when all Fe^2+^ has been used. It has therefore been suggested that fungi secrete secondary metabolites such as hydroquinones to reduce Fe^3+^ to Fe^2+^ (30). In spite of the decomposition potential of the Fenton reaction, it remains a major challenge for any living tissues to be exposed to concentrated doses of free radicals. Numerous adaptations have in fact evolved to remove low concentrations of free radicals from living tissues (31) so we conjectured that attine ants should have found ways to avoid such damage in case they would use Fenton chemistry.

Against this background we drew up a set of hypothetical mechanisms that might allow the leaf-cutting ant symbiosis to use Fenton chemistry, to sustain and control that reaction according to need, and to have it take place in a spatial setting where collateral damage to living tissues would be avoided. As it appears, the spatial issue appears to have been resolved because *Acromyrmex* colonies concentrate their chewed-up leaf-pulp in pellets of ca. 3 mm diameter at the top of their fungus gardens for some hours before that material is dispersed across the actively growing ridges of the fungal mycelium mass. However, if these balls would be the optimal location for a sustained Fenton reaction, the symbiosis would have been required to also evolve a fecal fluid enzyme that reduces Fe^3+^ while decomposing degradation products from the plant substrate. We hypothesized that glucose dehydrogenase (FP145), the most abundant ant encoded fecal fluid protein (Supplementary Table S1), could have this function using glucose as electron donor to reduce Fe^2+^, either directly or indirectly via an intermediate redox compound (Figure 2). We set up a number of biochemical assays to test this hypothetical scenario and report the results below.

**Figure 2.**
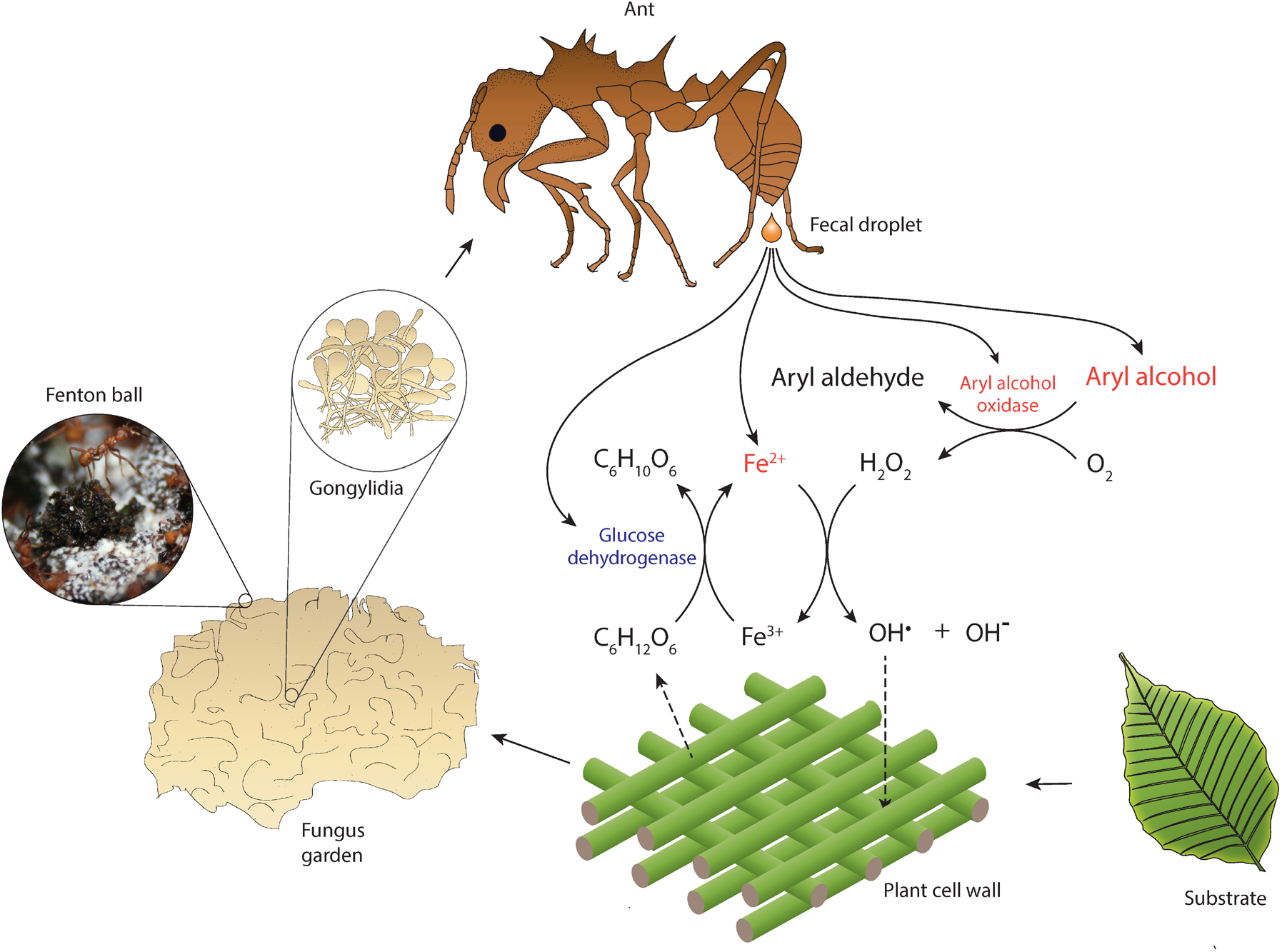
Hypothesized reaction scheme for the generation of hydroxyl radicals when ant fecal fluid interacts with ant-pretreated leaf pulp in temporary substrate pellets, based on the presences of enzymes in the total fecal fluid proteome (Supplementary Table S1). Fungal enzymes produced in gongylidia of the symbiotic garden-cultivar that the ants ingest pass unharmed through the gut to end up in the fecal fluid (17, 21, 23–25). After droplets of fecal fluid are deposited and become exposed to oxygen, the fungal oxidoreductases produce hydrogen peroxide when aryl alcohols are converted to aryl aldehydes. The hydrogen peroxide then reacts with reduced iron (Fe^2+^) to produce hydroxyl radicals (OH^•^) in a Fenton reaction, which aggressively breaks down cell-walls of the plant substrate. Oxidized iron (Fe^3+^) can then be reduced again by ant-encoded glucose hydrogenase, using glucose released via plant cell wall decomposition as electron donor. The leaf pulp substrate is initially concentrated green pellets of ca. 3 mm diameter distributed across the top of fungus gardens, which turn black in a few hours when subjected to Fenton-mediated degradation (inset image).

### Hydrogen peroxide production by the GMC oxidoreductases in ant fecal fluid

A phylogenetic tree of all five fungal GMC oxidoreductases isolated from fecal fluid, together with representative other such reductases from various basidiomycete fungi, showed that these five GMC oxidoreductases clustered within the aryl alcohol oxidases (Fig 3), while being distinct from the four other known functional groups (glucose oxidases, methanol oxidases, pyranose 2-oxidases and cellulose dehydrogenases) (32). We subsequently measured the amount of hydrogen peroxide that these GMC oxidoreductases produce with and without substrates, which showed that fecal fluid contained a substantial amount of hydrogen peroxide. Addition of the aromatic alcohol veratryl alcohol further increased this amount, whereas glucose and methanol did not have any effect (Fig 4A). While this confirmed the functional presence of aryl alcohol oxidases in the fecal fluid as hydrogen peroxide producer, we added a control assay to check that the increase in absorbance was directly caused by hydrogen peroxide by adding the enzyme catalase, which is a very effective and specific hydrogen peroxide degrading enzyme. As expected, catalase completely removed the signal, confirming that hydrogen peroxide is indeed naturally produced in fecal fluid (Fig 4A). Two other fecal fluid oxidoreductases (FP81 and FP186) were annotated as copper radical oxidases with close similarity to glyoxal oxidases. These enzymes may also produce hydrogen peroxide, using the small organic compound glyoxal and other aldehydes as substrate (33). Experimental addition of glyoxal to fecal fluid of *Acromyrmex* worker did indeed increase the amount of hydrogen peroxide, confirming that glyoxal oxidases are also present in the fecal fluid (Fig 4A).

**Figure 3.**
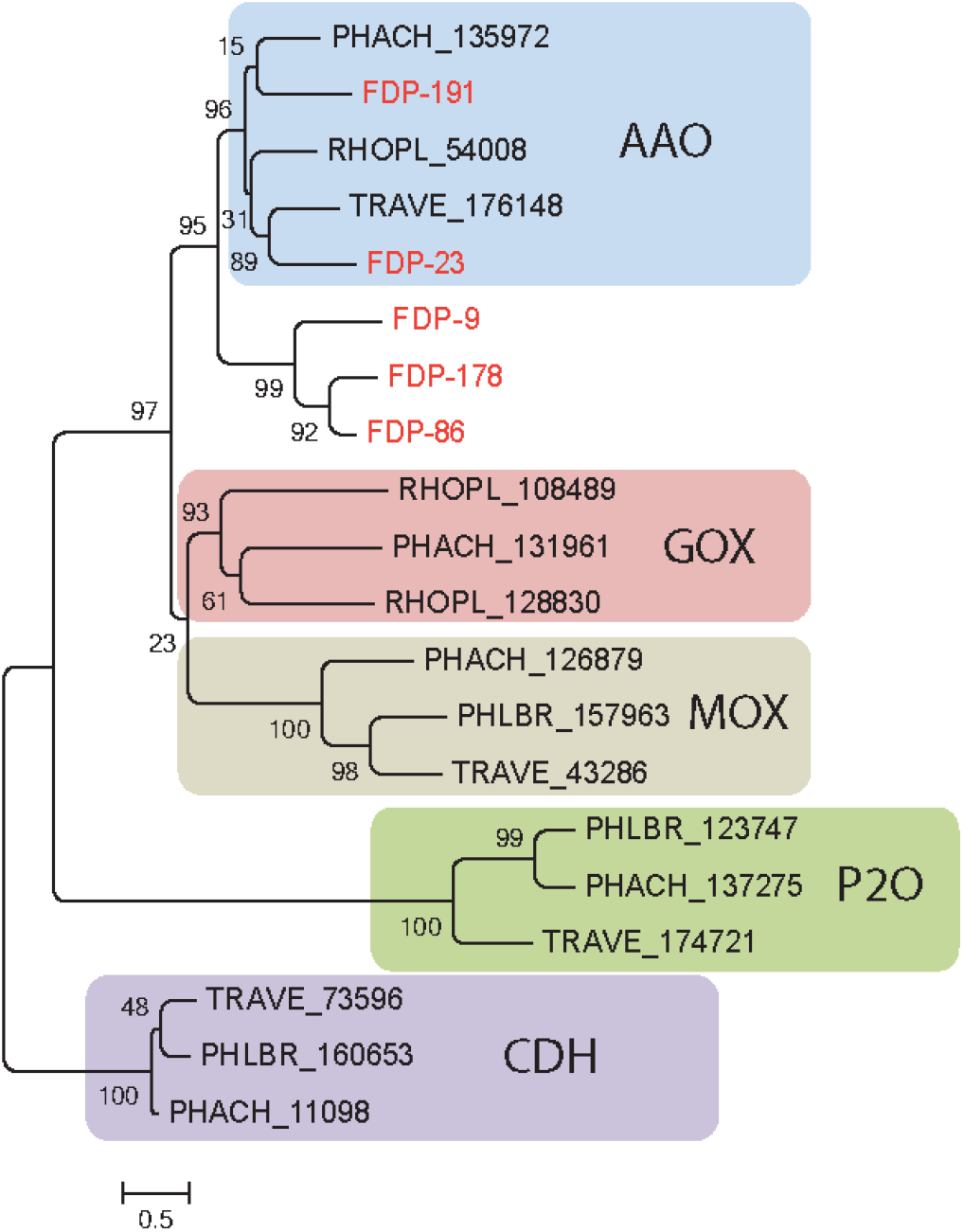
Gene tree of the five GMC Oxidoreductases identified in the ant fecal fluid (present study) and in representative other basidomycete fungi, all assigned to functional groups based on a previous study (32). All GMC Oxidoreductases from the fecal fluid (FP9, FP23, FP86, FP178 and FP191; red text) clustered among the known aryl-alcohol oxidases (AAO). Other closely related functional groups are glucose oxidases (GOX), methanol oxidases (MOX), pyranose-2 oxidases (P2O), and cellulose dehydrogenases (CDH), which were retrieved in *Phanerochaete chrysosporium* (PHACH), *Rhodonia placenta* (RHOPL), *Trametes versicolor* (TRAVE) and *Phlebia brevispora* (PHLBR), but not in fecal fluid of the leaf-cutting ant *A. echinatior*. Numbers are aLRT SH-like support values for nodes. The scale bar represents 0.5 substitutions per site.

**Figure 4.**
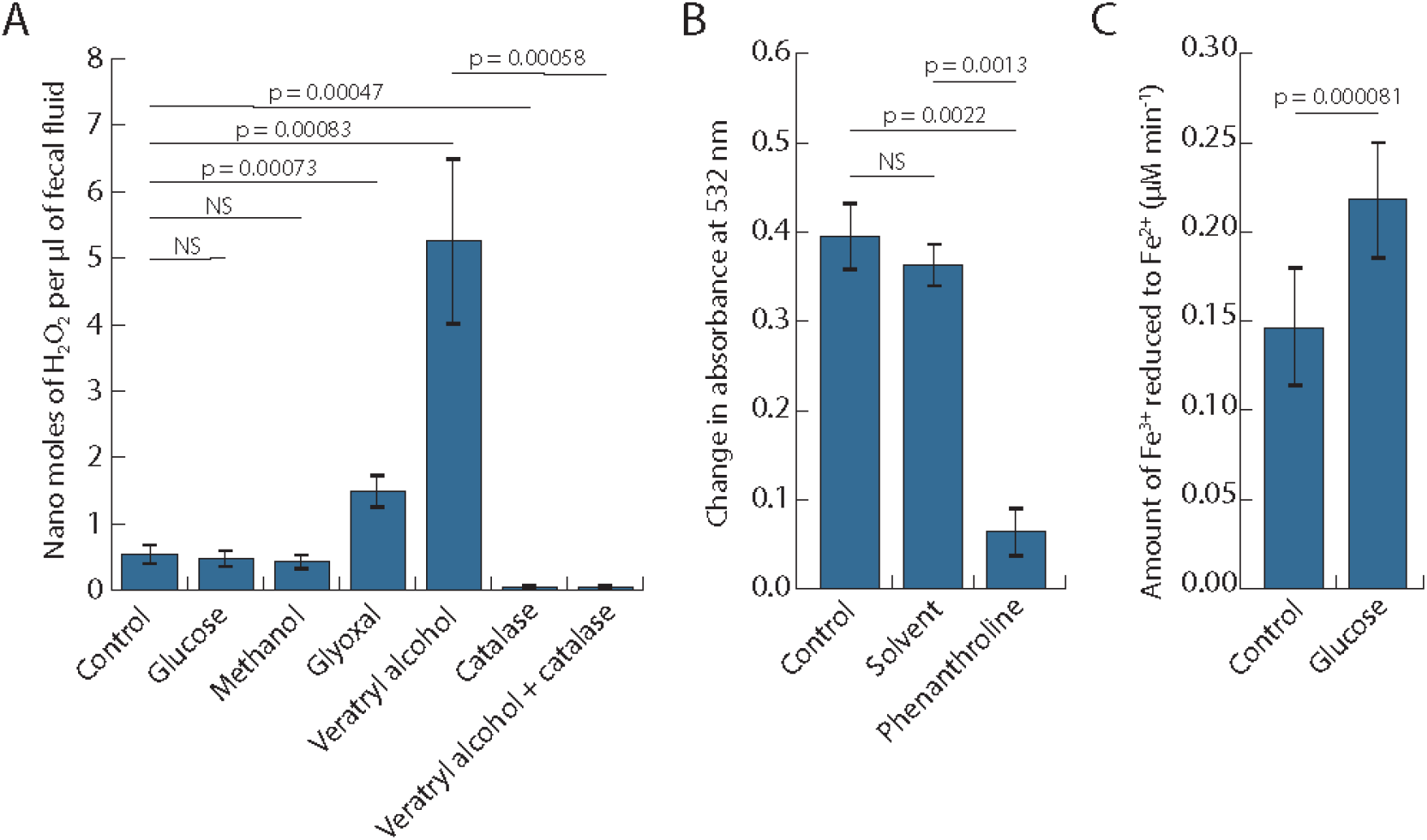
Bioassays to demonstrate that inorganic Fenton chemistry must be taking place when ant fecal fluid is exposed to oxygen while being deposited on leaf pulp pellets munched by the ants, testing the key interactions hypothesized in Figure 2. (**A**) Bar plot showing the concentrations (means ± SE across 6 colonies) of hydrogen peroxide in fecal fluid after adding potential substrates (glucose, methanol, glyoxal or veratryl alcohol) of GMC oxidoreductases with or without the hydrogen peroxide degrading enzyme catalase. One-way ANOVA showed a highly significant overall effect of treatments: *F*_6,36_ = 14.597, p = 2.863e-08, and pairwise posthoc t-tests on matching samples from the same ant colonies (corrected for multiple testing with the Holm-Bonferroni method) confirmed both the enhancing effects of AAOs and the inhibiting effect of catalase (p-values in plot unless non-significant (NS). (**B**) Deoxyribose assays showing that ant fecal fluid has the capacity to produce hydroxyl radicals (means ± SE across 6 colonies). Phenanthroline is known to work as an iron chelator and significantly reduced degradation of 2-Deoxy-D-ribose while the solvent (methanol) of phenanthroline did not. Paired t-tests followed the same protocol as in the A-panel except that the overall ANOVA was omitted because there were only three means to compare. (**C**) Ferrozine assay (means ± SE across 6 colonies) showing the capacity of ant fecal fluid to reduce Fe^3+^ to Fe^2+^, confirming that addition of glucose increases the rate of iron reduction. Statistics as in the B-panel.

### Does ant fecal fluid contain sufficient iron to maintain Fenton chemistry?

For the Fenton reaction to take place, the produced hydrogen peroxide must react with Fe^2+^ atoms. Brown-rot fungi are believed to rely on iron in their wood substrate (34) but the iron that may be present in the leaf material collected by ant workers is not yet available when this plant biomass has only just been chewed-up by the ants. Given that 85% of the fecal fluid enzymes are derived from the ant-ingested fungal food, we therefore investigated whether the fecal fluid might provide the required iron. Making fungal iron available would be adaptive for the ant farming symbiosis as a whole because direct use of iron from live plant tissues might well be constrained by plants also producing iron chelator compounds to inhibit the growth of plant pathogens (35). We measured the iron levels in fecal fluid, and found an average (± SD) level of 370 ± 110 μM iron across 6 ant colonies). This is a very high concentration if one takes into account that human blood contains about 18 μM iron (36), i.e. less than 5% of the iron concentration in ant fecal fluid. It thus appears likely that fecal fluid is instrumental in maintaining the Fenton reaction and that the ants control this process by actively regulating the volume of fecal fluid to be added to the balls of freshly munched leaf pulp.

### Are hydroxyl radicals Fenton produced and can the ants control iron reduction?

The hydroxyl radicals that Fenton chemistry should produce can be measured via their ability to break down 2-Deoxy-D-ribose into thiobarbituric acid-reactive substances (TBARS) (37). To quantify this affect, we adapted the standard 2-Deoxy-D-ribose assay to accommodate the small volumes of fecal fluid that could practically be collected directly from the ants and found pronounced amounts of hydroxyl radicals to be present in the ant fecal fluid (Fig 4B). Adding a strong metal chelator 1,10-phenanthroline caused a significant reduction in absorbance, consistent with the hydroxyl radicals being produced by Fenton chemistry, which should be inhibited by 1,10-phenanthroline sequestering iron atoms (38). This effect was not observed when only the solvent of the 1,10-phenanthroline chemical (methanol) was added.

The Fenton reaction can only proceed as long as there are Fe^2+^ atoms present to be converted to Fe^3+^ atoms. Brown-rot fungi are believed to maintain their Fenton reactions via enzyme systems that convert Fe^3+^ back to Fe^2+^ (29, 34), but leaf-cutting ants pre-digest chewed-up leaf pulp with fecal fluid before it is being made available to the fungal mycelium of their garden symbiont, so we tested the hypothesis that ant encoded glucose dehydrogenase mediates the reduction of Fe^3+^ to Fe^2+^ using glucose as electron donor. We indeed found a considerable capacity for Fe^3+^ reduction in fecal fluid, which was further increased by the addition of glucose. This was consistent with Fe^3+^ being actively recycled into Fe^2+^ via fecal fluid compounds and with glucose dehydrogenase most likely being responsible for at least part if not all of this activity (Fig 4C).

## Discussion

### *Acromyrmex* fecal fluid has a highly specialized proteome

The presence of fungal enzymes in the fecal fluid of leaf-cutting ants was suggested already in the 1970’s based on the biochemical similarity of fecal fluid and fungal proteases (17, 18). We achieved increasing confirmation of this idea in a series of previous studies (21, 23–25), but our present study presents the first overall proteome that is complete enough to allow inferences of hitherto unknown chemical processes. The comprehensive list of *A. echinatior* fecal fluid proteins that we obtained (Supplementary Table S1) was now also sufficiently replicated to confirm that the large majority of the identified proteins are fungal in origin and serve the farming symbiosis after being deposited on new plant substrate via the ant fecal fluid. When run on SDS-Page gels (23), the fecal fluid proteins always gave the same banding pattern, rejecting the null hypothesis that they are a random collection of proteins that happen to escape digestion. Overall, 101 of the 174 identified proteins were found in at least three samples originating from three different ant colonies, and for many of them we were able to measure the corresponding enzymatic activity in fecal fluid using biochemical assays (this study and (21, 23–25)), consistent with predicted protein functions as fresh leaf decomposition agents. These results strongly suggest that most if not all of the fecal proteins have obligate condition-dependent adaptive significance, have been selected to avoid degradation in the digestive system of the ants, and serve the efficiency of the entire symbiosis between farming ants and fungal cultivars.

### Multiple lines of evidence for fecal fluid Fenton chemistry after mixing with fresh leaf pulp

The key compounds hypothesized to be involved in inorganic Fenton chemistry (GMC oxidoreductases, hydrogen peroxide, high concentrations of iron, glucose dehydrogenase) were always found in all of the colonies for which we investigated fecal fluid samples. Compared to white rot fungi, brown rot fungi are believed to have a reduced array of enzymes targeting lignocellulose, and to specifically lack lignin peroxidases, which is also the case for *L. gongylophorus* (16). Brown rot fungi compensate this lack of enzymatic potential by using Fenton chemistry to degrade lignocellulose while producing extracellular hydroxyl radicals (39). In that Fenton reaction, hydrogen peroxide reacts with ferrous iron (Fe^2+^) and hydrogen ions to produce ferric iron (Fe^3+^), water and hydroxyl radicals (29, 34). The very reactive hydroxyl radicals produced in these free-living fungi are known to damage a wide range of biomolecules (37). Their putative functions are to release soluble carbohydrate fragments from plant cell walls (40) and to reduce the viscosity of solutions of plant cell wall components such as xyloglucan, pectin and cellulose (41) as expected when degradation of these polysaccharides has taken place. The small size of hydroxyl radical molecules allows them to initiate penetration and cleavage of lignin, cellulose and hemicellulose polymers as precursors for further degradation into assimilatory monomers by specific enzymes (34). Brown rot fungi are believed to use GMC oxidoreductases or copper radical oxidases to aerobically oxidize fungal alcohols or aldehydes in order to produce hydrogen peroxide, while obtaining the required iron from the wood substrate (34). It has been suggested that fungal laccases are then oxidizing hydroquinones into the semiquinones that react with ferric iron (Fe^3+^) to produce quinones and ferrous iron (Fe^2+^), which would produce the recycling process needed to maintain the Fenton reaction over time (29).

It is intriguing that the details of Fenton chemistry in brown rot fungi have remained rather little known in spite of the economic importance of these fungi (39), and that the leaf-cutting ants offered an unexpected window for understanding this elusive process. The laccases, GMC oxidoreductases and copper radical oxidases were similarly found in high quantities in the fecal fluid of *Acromyrmex* leaf-cutting ants, consistent with Fenton chemistry being a standard procedure in the attine fungus-farming symbiosis to degrade plant cell walls and/or toxic secondary plant polymers meant to deter herbivores (Fig 2). We could verify that the identified GMC oxidoreductases were aryl alcohol oxidases (Fig 3), which are known to produce hydrogen peroxide from aromatic alcohols. We also verified that fecal fluid contains concentrated hydrogen peroxide and that the concentration increased further after adding veratryl alcohol or glyoxal, which are substrates for GMC oxidoreductases and copper radical oxidases, respectively (33, 42). We further documented that the iron content of the fecal fluid was surprisingly high (ca. 20 times higher than in human blood), suggesting that the velocity of Fenton chemistry is not limited by metal ions. Also the acid pH of ca. 4 in the fecal fluid should be ideal for Fenton chemistry (43, 44). Finally, our deoxyribose assay showed that fecal fluid components degrade deoxyribose into thiobarbituric acid-active products, which can only be explained by the presence of hydroxyl radicals (37). To obtain more direct evidence for the produced hydroxyl radicals contributing to plant substrate breakdown, future research should focus on using Fourier transform infrared spectroscopy to analyze plant cell wall preparations subjected to fecal fluid degradation, a type of analytical chemistry that was beyond the scope of the present study. Without such validation, we cannot rule out other possible functions of these radicals such as general sanitation of the plant substrate. However, such a function has never been conjectured or shown for the ant fecal fluid, whereas brown rot fungi are widely believed to use hydroxyl radicals for substrate degradation (39) and specialized functions of the ant fecal fluid for plant substrate degradation are uncontroversial (17–25). Our Fenton-chemistry interpretation is therefore supported by consistent inferential evidence, but in need of further experimental validation.

The high concentrations of laccase LgLcc1 (FP097) in the ant fecal fluid suggest a parallel mechanism of maintaining reliable conversion of Fe^3+^ back to Fe^2+^ to the one known from brown rot fungi (29). However, the glucose dehydrogenase (FP145) encoded by the ant genome was also abundant in the fecal proteome and could convert Fe^3+^ back to Fe^2+^ by simultaneous oxidization of glucose. We were also able to verify this function with a Ferrozine based assay (45), including its dependence on glucose availability. Interestingly, however, preliminary experiments indicated that adding laccase substrates did not influence the velocity of this reaction, which suggests that glucose liberated from the plant polysaccharides is primarily used to keep the Fenton reaction going. This substrate dependence may provide an additional feedback loop for the farming symbiosis to control the inorganic Fenton chemistry process. The ant-derived glucose dehydrogenase (FP145) alternative might also allow the ants to use the fungal laccases specifically for the decomposition of secondary plant compounds as found in one of our previous studies (24). However, that study also suggested that LgLcc1 (FP097) came under positive selection in the common ancestor of the leaf-cutting ants to avoid digestion in the ant digestive tract, which would be consistent with the feedback loop redundancy explanation.

### Symbiotic complementarity is necessary for making fast Fenton chemistry work

The assembly of complementary components from different sources implies that the fungal cultivar and the farming ants synergistically cooperate to accomplish the continued production of hydroxyl radicals, but only as long as Fenton chemistry is needed. The ants do not appear to keep adding fecal fluid to existing Fenton pellets and they take them apart when they have turned black after some hours to distribute the Fenton exposed leaf pulp over the actively growing ridges of the fungus garden where fungal decomposition takes over. Both the processing efficiency and the ultimate behavioral control by the ants strongly suggest refined co-adaptation at the molecular level to enable colonies to operate as functional herbivores. An earlier study found a high expression of an NADH-quinone oxidoreductase in fungus gardens of *A. echinatior* (46), an enzyme converting quinones to hydroquinones that can react with Fe^3+^ to form Fe^2+^. This is consistent with the ant encoded glucose dehydrogenase being relevant only for the degradation process in the Fenton pellets and with NADH-quinone oxidoreductases taking over after the leaf pulp has been dispersed over the garden and the fungal hyphae decompose what remains of the substrate.

Aryl alcohol oxidases use aromatic alcohols as a substrate to produce hydrogen peroxide. Free-living fungi commonly produce aromatic veratryl alcohol as a secondary metabolite, as for example in white rot fungi that decompose lignin with peroxidase enzymes (47) rather than with Fenton chemistry as brown rot fungi do. Veratryl alcohol is derived from phenylalanine via the phenylalanine/tyrosine pathway that has previously been shown to be upregulated in the gongylidia of the *A. echinatior* cultivar *L. gongylophorus* (48). This once more underlines the co-adapted utility of such compounds being ingested but not digested by the ants so they can function in fecal fluid to facilitate the initial stages of substrate degradation. The hydrogen peroxide needed in the Fenton reaction may also be derived from the activity of the glyoxal oxidases that we identified, which use various aldehydes as substrate for hydrogen peroxide production (33). Such aldehydes could then either originate from the oxidized aryl alcohols produced by the aryl alcohol oxidases, or could be direct degradation products from plant cell wall components (33). These multiple parallel pathways suggest that the co-evolved interactions between inorganic Fenton chemistry and organic enzyme functions are likely to be operational under a variety of environmental (i.e. substrate) conditions and thus highly robust.

Finally, as we explained in the introduction, it is important to note that hydroxyl radicals are highly toxic also for the organisms that producing them. This probably constrains the use of naturally produced hydroxyl radicals for biomass degradation unless the Fenton chemistry producing these radicals remains distal to hyphal growth and is terminated by the time hyphae grow into the substrate patches pre-treated in this manner. This complication will slow down the rate of decomposition in free-living fungi, but the leaf-cutting ant symbiosis appears to have resolved this constraint by compartmentalizing the complementary elements of first aggressive substrate break down. Although the necessary compounds for the Fenton reaction are all present in the guts of the ants, the low oxygen pressure there would presumably prevent the production of hydrogen peroxide inside the ants and thus any collateral damage to ant tissues. Defecation onto concentrated pellets of chewed-up leaf pulp thus ensures that the reaction does not start until atmospheric oxygen is available. The Fenton pellets thus appear to function as small bioconversion reactors that are spatially isolated from both the fungal and the ant tissues so that potentially harmful side-effects are avoided. Also in this respect, robustness against potential malfunction appears to have evolved. Our recent genome sequencing study of *Acromyrmex* ants and their cultivars revealed that *A. echinatior* has more Glutathione S-transferase genes than other (non-attine) ants, despite having fewer detoxification genes in general (49). Glutathione S-transferase is known to be involved in the breakdown of oxidized lipids, which would likely be produced if reactive oxygen species would be formed in the gut and react with the lipid cell membranes of the gut tissues. This suggests that physiological mechanisms to ameliorate inadvertent damage caused by hydroxyl radicals may be in place should they appear prematurely in the ant gut.

### Implications for our general understanding of the leaf-cutting ant symbiosis

The extent to which cellulose is degraded in the fungus garden has been widely disputed (see (46) and references therein). It is noteworthy that the fecal fluid proteome only contained a single 1,4-beta-glucanase (FP55) to target the cellulose backbone, and that this protein was found in low quantities and in only three of the four examined colonies. We also found a single 1,4-beta-glucosidase (FP73) that releases glucose moieties from cellobiose and oligosaccharides, but this enzyme is also targeting hemicellulose in addition to cellulose. Finally, effective degradation of cellulose normally requires a cellobiohydrolase enzyme, which was only found in one colony in very low quantities (Supplementary Table S1). These findings are consistent with our earlier inference (46, 50) that cellulose degradation is not a primary function of the fecal fluid, supporting earlier findings that recalcitrant plant cell wall cellulose is not degraded to a significant degree in *Acromyrmex* fungus garden, but discarded from the colony as waste (50). Our finding that Fenton chemistry may be used for breaking chemical bonds in the cell wall lignocellulose and likely sufficient to give hyphal degradation enzymes access to their target substrates in the interior of plant cells appears to corroborate this notion. More complete cellulose degradation would release an excess of assimilatory sugars for the fungal symbiont and without simultaneously providing nitrogen or phosphorus, which are the limiting factors for growth in any functional herbivore (51). Such sugars could well be a burden for the symbiosis if they would allow inadvertent microorganisms to thrive in fungus gardens (46).

While the documentation of Fenton chemistry substantially increases our understanding of the way in which *Acromyrmex* leaf-cutting ants could become crop pests by robust functional herbivory, it is intriguing that we have never observed Fenton pellets in any of the *Atta* colonies that we have maintained in the lab for ca. 25 years. This suggests that the colonies of this other genus of leaf-cutting ants, which are ca. two orders of magnitude larger in size and considerably more damaging as agricultural pests have evolved alternative mechanisms to boost the efficiency and robustness of their functional herbivory. Given that also *Atta* colonies produce enormous amounts of cellulose rich waste, we do not expect that the explanation for *Atta*’s success as functional herbivores will be fundamentally different because both genera rear very closely related lineages of the *L. gongylophorus* cultivars in sympatry at our Panamanian sampling site (52) and across their Latin American distributions (53). We suspect, therefore that obtaining a fecal fluid proteome of *Atta* might shed interesting light on whether these ants can also use forms of inorganic chemistry, or whether they rely on an enzymatic innovation not available in *Acromyrmex*. Another point of interest is that human *in vitro* experiments with Fenton chemistry to pretreat plant biomass for more efficient conversion into valuable biofuel products have produced up to fivefold increases in the production of sugars from feedstock, although the efficiency remains highly dependent on the plant material used (54–56). A major challenge in these human bioconversion applications has been that the Fenton generated reactive oxygen species degrade the very enzymes that are instrumental in realizing conversion efficiency. An interesting venue for further research is therefore to elucidate how *Acromyrmex* fecal fluid degradation enzymes avoid being harmed by the hydroxyl radicals that accompany them.

Finally, we are used to think of our own crops as being in continuous need of manure to boost nutrient availability and growth. The term manure now seems inappropriate for attine ant fecal fluid even though deposition of fecal droplets in the fungus garden seems to suggest this analogy. In retrospect, abandoning this term seems logical, because the ants rear a heterotrophic crop and it seems hard to imagine how fresh leaf substrate can be further enriched with nutrients. However, our present results suggest that it is also ambiguous to use the term ‘fecal’ for the droplets that leave the hindguts of leaf-cutting ants. In a structural sense this is fecal material, but in a functional sense this fluid now seems to be analogous to a joint symbiotic ‘bloodstream’ with the primary function of optimizing the distribution of gongylidia-produced decomposition compounds to simultaneously maintain the compound superorganismal body of ants, ant brood and fungus garden. In this perspective, one might even argue that it has become ambiguous which partner is the farmer or the crop, because the relationship is fully symmetrical and the cultivar has remarkable agency even though it neither has brains to coordinate or legs to move around.

## Methods

### Biological material

Colonies of *Acromyrmex echinatior* (numbers Ae150, Ae160, Ae263, Ae322, Ae356, Ae372, Ae480 and Ae490) were collected in Gamboa, Panama between 2001 and 2010 and maintained in the laboratory at ca. 25°C and ca. 70% relative humidity (57) where they were supplied with a diet of bramble leaves, occasional dry rice, and pieces of apple. Fecal droplets were collected by squeezing large worker ants with forceps on the head and abdomen until they deposited a drop of fecal fluid. For mass spectrometry, 0.5 μl of double distilled and autoclaved water was added to the droplet before it was collected with a micro pipette and transferred to an Eppendorf tube with Run Blue loading buffer (Expedeon Inc.). For the iron assay, Fenton reaction assay, hydrogen peroxide assay and iron reduction assays, the fecal droplets were deposited on parafilm and collected undiluted with a 5 μl glass capillary.

### Mass spectrometry

Ten fecal droplets from each of the four colonies were run on a 12% Run Blue polyacrylamide gel (Expedeon Inc.) in a Mini-Protean II electrophoresis system (Biorad) for about 10 min at 90 mA. The gel was stained in Instant Blue Coomassie (Expedeon Inc.) over night at room temperature. The stained part of each lane was then cut out with a scalpel and transferred to Eppendorf tubes, after which the samples were sent to Alphalyse (Denmark) for analysis.

The protein samples were reduced and alkylated with iodoacetamide (i.e. carbamidomethylated) and subsequently digested with trypsin. The resulting peptides were concentrated by Speed Vac lyophilization and redissolved for injection on a Dionex nano-LC system and MS/MS analysis on a Bruker Maxis Impact QTOF instrument. The obtained MS/MS spectra were analysed using the MaxQuant software package version 1.5.4.1 (58) integrated with the Andromeda search engine (59) using standard parameters. The spectra were matched to a reference database of the predicted *Acromyrmex echinatior* proteome (49) combined with published predicted *Leucocoprinus gongylophorus* proteomes originating from the *Acromyrmex echinatior*, *Atta cephalotes* and *Atta colombica* fungal symbionts (16, 49, 60), as well as single protein sequences predicted from manually PCR amplified and sequenced cDNA sequences (21, 23–25, 61). As these published fungal proteomes contained many redundant protein sequences, a non-redundant reference protein database was constructed using CD-Hit (62) keeping only the longest sequences of the sets of 100% matching sequences. Annotation of the identified protein sequences was performed using Blast2GO (33) combined with manual BlastP searches against the NCBI nr database.

### Iron level measurements

Fecal fluid iron content was measured according to Fish 1988 (63). Fecal fluid from 20 workers from each of six colonies (Ae150, Ae160, Ae263, Ae356, Ae480, Ae490) was collected with glass capillaries and weighed in 0.2 ml PCR tubes to determine the volume of each sample, assuming a density of 1 g/ml. The samples were diluted to 100 μl with 0.1 M HCl, mixed with Reagent A (0.142 M KMnO4, 0.6 M HCl), and incubated for 2 hours at 60°C. Subsequently 10 μl Reagent B (6.5 mM Ferrozine, 13.1 mM neocuproine, 2 M ascorbic acid, 5 M ammonium acetate) was added to each sample and incubated for 30 minutes at room temperature, after which 150 μl of this mixture was transferred to a microtiter plate and the absorbance at 562 nm measured in an Epoch microplate spectrophotometer (BioTek). Samples with known amounts of iron were processed in parallel with the unknown samples to produce a standard curve from which the iron content of the unknown samples could be determined. Three series of standards were made from ferrous ethylenediammonium sulfate tetrahydrate, ferrous sulfate heptahydrate, and ferric chloride hexahydrate. In all cases a solution of 5 mg/l iron in 0.1 M HCl was used to make a dilution series ranging from 78.1 μg/l to 5 mg/l. In addition, a blank sample without iron was included and used to correct for background absorbance.

### Hydroxyl radical measurements

0.5 μl fecal fluid from each of the colonies Ae150, Ae160, Ae263, Ae356, Ae480, Ae490 was diluted with 0.5 μl distilled H_2_O and mixed with either 1 μl H_2_O, 1 μl 40 mM 1,10-phenanthroline (dissolved in methanol to 200 mM and diluted 1:4 with distilled H_2_O) or 1 μl 20% methanol. After 15 minutes incubation at room temperature, 2 μl substrate (100 mM 2-Deoxy-D-ribose in 0.1 M Sodium acetate buffer pH 5.0) was added, and samples were incubated for 30 minutes at room temperature. Then, 4 μl of a solution of 1% 2-Thiobarbituric acid in 50 mM NaOH was added followed by addition of 4 μl of 2.8% Trichloroacetic acid. The samples were then incubated for 20 minutes at 99.9° C in a PCR machine, after which 2 μl was used for measuring the absorbance at 532 nm using a NanoDrop ND-1000 spectrophotometer (37). Control samples without addition of 2-Deoxy-D-ribose were made in parallel and the absorbance value subtracted from the absorbance values of the samples. Similarly, control samples with 2-Deoxy-D-ribose added but without fecal droplets were made in parallel so also their absorbance value could be subtracted from those of the samples. Control samples without both 2-Deoxy-D-ribose and fecal droplets were also run in parallel and these absorbance values were added to the final values to compensate for subtracting the absorbance values of two controls from each sample value.

### H_2_O_2_ measurements

0.5 μl fecal fluid from each of the colonies Ae150, Ae160, Ae263, Ae356, Ae480, Ae490 was mixed with either 150 μl demineralized H_2_O or with 150 μl 10 mM glucose, 10 mM glyoxal, 10 mM veratryl alcohol, 10 mM methanol or 100 U catalase. The samples were incubated for 60 minutes at 30° C in a PCR machine. To avoid direct oxidation of the amplex red substrate by laccases in the fecal fluid, the samples were transferred to Amicon Ultra-0.5 Centrifugal filters with 3 Kda membranes (Merck Millipore UFC500396) and centrifuged at 14,000 g for 30 minutes. 100 μl flow through was transferred to a microtiter plate and 100 μl amplex red reagent (100 μM Ampliflu red, 2 U/ml Horse Radish Peroxidase, in 0.1 M phosphate buffer pH 7.4) (64) was added, after which the absorbance at 560 nm was measured for 30 minutes using an Epoch microplate spectrophotometer (BioTek). This absorbance measure was then used to calculate the H_2_O_2_ concentration using a standard curve made from a twofold serial dilution of H_2_O_2_ ranging from 1.56 μM to 100 μM and including a blank sample without H_2_O_2_. The standard samples were assayed in parallel with the fecal fluid samples.

### Iron reduction measurements

One microliter fecal fluid from each of the colonies Ae150, Ae160, Ae263, Ae356, Ae480, Ae490 was mixed with 71 μl demineralized H_2_O, 10 μl 1 mM FeCl_3_ · 6 H_2_O, 115 μl 0.1 M Acetate buffer pH 4.4, and either 23 μl H_2_O or 23 μl 100 mM Glucose. Subsequently, 10 μl 1% FerroZine was added and the absorbance at 562 nM was measured during a 30 minutes period using an Epoch microplate spectrophotometer (BioTek) (45). The rates of change in absorbance from 5 to 30 minutes were used to calculate the amount of Fe^3+^ reduced to Fe^2+^ per minute, using a standard curve made from a twofold serial dilution of FeSO_4_ · NH_3_CH_2_CH_2_NH_3_SO_4_ · 4 H_2_O, ranging from 5 mM to 0.078 mM and including a blank sample without iron. The standard samples were assayed in parallel with the fecal fluid samples.

### Statistical data analyses

All analyses were done in RStudio Version 0.99.491. Shapiro-Wilk tests were used to test whether differences in response variables between treatments within the same ant colonies followed a normal distribution. For hydrogen peroxide measurements a square root transformation of response data was applied to make the data pass the Shapiro-Wilk test. Planned comparisons between treatments were performed with paired t-tests (by pairing measurements of samples coming from the same ant colonies). P-values were adjusted for multiple comparisons using the Holm-Bonferroni method.

### Phylogenetic analyses

The five identified fecal fluid GMC oxidoreductase sequences were aligned with 15 basidiomycete GMC oxidoreductase sequences belonging to each of the five functional GMC oxidoreductase protein groups presented in (32), using M-coffee [http://tcoffee.crg.cat/apps/tcoffee/do:mcoffee] with default parameter settings. The alignment file was then used in PhyML with Smart Model Selection (http://www.atgc-montpellier.fr/phyml-sms/) and aLRT SH-like support values. The best DNA substitution model turned out to be LG+G+I+F.

## Acknowledgements

We thank the Smithsonian Tropical Research Institute (STRI), Panama, for providing logistic help and facilities during our field work in Gamboa.

## Ethics and consent to participate

JJB obtained permission to collect ant colonies and export them from Panama to Denmark from the Autoridad Nacional del Ambiente y el Mar (ANAM), Panama.

## Authors’ contributions

Conceived and designed the experiments: MS, JJB. Performed the experiments: MS. Analyzed the data: MS. Wrote the paper: MS, JJB.

## Availability of data and materials

The mass spectrometry data generated in this project as well as the amino acid sequences used for protein identification have been submitted to the PRIDE Archive with accession number PXD016395. [<u>During the review process the data can be accessed with the Username</u>: <u>and the Password: orlQ6EOU</u>]

## Supplementary files

Supplementary Table S1: Data for all fecal fluid proteins identified in the study.

